# Physics-based predictive simulations to explore the differential effects of motor control and musculoskeletal deficits on gait dysfunction in cerebral palsy: a retrospective case study

**DOI:** 10.1101/769042

**Authors:** Antoine Falisse, Lorenzo Pitto, Hans Kainz, Hoa Hoang, Mariska Wesseling, Sam Van Rossom, Eirini Papageorgiou, Lynn Bar-On, Ann Hallemans, Kaat Desloovere, Guy Molenaers, Anja Van Campenhout, Friedl De Groote, Ilse Jonkers

## Abstract

Model-based simulations of walking have the theoretical potential to support clinical decision making by predicting the functional outcome of treatments in terms of walking performance. Yet before using such simulations in clinical practice, their ability to identify the main treatment targets in specific patients needs to be demonstrated. In this study, we generated predictive simulations of walking with a medical imaging based neuro-musculoskeletal model of a child with cerebral palsy presenting crouch gait. We explored the influence of altered muscle-tendon properties, reduced neuromuscular control complexity, and spasticity on gait function in terms of joint kinematics, kinetics, muscle activity, and metabolic cost of transport. We modeled altered muscle-tendon properties by personalizing Hill-type muscle-tendon parameters based on data collected during functional movements, simpler neuromuscular control by reducing the number of independent muscle synergies, and spasticity through delayed muscle activity feedback from muscle force and force rate. Our simulations revealed that, in the presence of aberrant musculoskeletal geometries, altered muscle-tendon properties rather than reduced neuromuscular control complexity and spasticity were the primary cause of the crouch gait pattern observed for this child, which is in agreement with the clinical examination. These results suggest that muscle-tendon properties should be the primary target of interventions aiming to restore a more upright gait pattern for this child. This suggestion is in line with the gait analysis following muscle-tendon property and bone deformity corrections. The ability of our simulations to distinguish the contribution of different impairments on walking performance opens the door for identifying targeted treatment strategies with the aim of designing optimized interventions for neuro-musculoskeletal disorders.

## 1 INTRODUCTION

Cerebral palsy (CP) is the most common cause of motor disability amongst children, affecting 2 to 3 per 1000 live births in Europe (Surveillance of Cerebral Palsy in Europe (2002)). CP is caused by a non-progressive lesion in the immature brain that may induce inabilities to selectively control muscles, spasticity, and weakness. These deficits undermine walking performance and, over time, lead to secondary impairments, such as bone deformities and muscle contracture, that may further deteriorate walking abilities (Gage et al. (2009)). Numerous treatments target these impairments with the aim of improving walking performance, such as single-event multi-level orthopedic surgeries (SEMLS) to correct multiple bone and muscle impairments in a single intervention (McGinley et al. (2012)). Yet walking involves complex interactions between the musculoskeletal and motor control systems, which are both impaired in CP. Hence, the treatment outcome does not only depend on the success of the intervention in terms of musculoskeletal remediation but also on the remaining motor control (Schwartz et al. (2016)). As a result, over the last decades, only modest, unpredictable, and stagnant treatment outcomes have been documented for children with CP (Schwartz (2018)). For example, SEMLS have been reported to improve walking performance in only 25 to 43% of the patients (Filho et al. (2008); Chang et al. (2006)) and to lead to clinically meaningful improvements over natural progression in only 37% of the cases (Rajagopal et al. (2018)). Computer models that can predict the functional outcome of treatments on walking performance have the potential to improve this success rate by allowing clinicians to optimize the clinical decision making (e.g., by discriminating the effects of musculoskeletal restoration due to surgical interventions to those from tone reduction and physical therapy targeting motor control impairments). However, predictive simulations are not yet applied in clinical practice, in part due to computational and modeling challenges.

Predictive simulations generate novel movements based on a mathematical model of the neuro-musculoskeletal system without relying on experimental data. Typically, these simulations consist in identifying muscle excitations that follow a certain control strategy and drive the musculoskeletal model to achieve a movement-related goal (e.g., moving forward at a given speed). For such simulations to be valuable in predicting the functional outcome of treatments on walking performance, they should be based on models that are complex enough to describe the musculoskeletal structures and motor control processes underlying walking that may be impaired and thus affected by treatment. Yet these complex models are computationally expensive in predictive simulations (Anderson and Pandy (2001); Miller (2014); Song and Geyer (2015); Lin et al. (2018)) and, therefore, their ability to predict the variety of gaits encountered under different conditions (e.g., healthy and pathological gaits) has been only scarcely explored in the literature. We recently developed a simulation framework to generate rapid (i.e., about 30 minutes of computational time) predictive simulations of gait with complex models (Falisse et al. (2019)). Further, we demonstrated the ability of our framework to predict the mechanics and energetics of a broad range of gaits, suggesting that our models and simulations were sufficiently generalizable for use in clinical applications. Nevertheless, the ability of our simulations to identify the main treatment targets in specific patients remains untested. Specifically, for children with CP, simulations should allow distinguishing the effects of musculoskeletal versus motor control impairments on walking performance to be able to help clinicians optimize treatments.

Predicting the effects of impairments on walking performance in children with CP requires that the neuro-musculoskeletal model captures these impairments. In this work, we focus on two types of impairments: motor control impairments that include spasticity and non-selective muscle control, and musculoskeletal impairments that include bone deformities and altered muscle-tendon properties.

Spasticity has been described as a velocity-dependent increase in tonic stretch reflex responses resulting from hyper-excitability of the stretch reflex (Lance (1980)). Following such description, models have been developed to describe the measured response in muscle activity (i.e., electromyography (EMG)) to passive stretches based on feedback from muscle velocity (van der Krogt et al. (2016)). However, we previously showed that a model based on feedback from muscle force and force rate better explains the response of spastic hamstrings and gastrocnemii than length- and velocity-based models (Falisse et al. (2018)). Further, we found that a force-based model could predict muscle activity in agreement with pathological EMG during gait. While spasticity manifests during passive stretches, its influence during gait remains unclear (Dietz and Sinkjaer (2007)). Incorporating spasticity models in predictive simulations would allow evaluating the impact of spasticity on gait performance, providing insights into the role of spasticity during gait. Further, modeling spasticity is a prerequisite for simulating the effects of treatments aiming to reduce spasticity, such as botulinum toxin type A (BTX-A) injections.

The inability to selectively control muscles has been described through muscle synergies (Ivanenko et al. (2004)), which are independent groups of muscles activated in a fixed ratio by a single input signal. Children with CP have been shown to use fewer synergies (i.e., a simpler neuromuscular control strategy) than typically developing (TD) individuals during walking (Steele et al. (2015)) as well as to use synergies exhibiting a greater stride-to-stride variability (Kim et al. (2018)). However, assessing the relationship between simpler neuromuscular control and impaired gait is difficult. For example, Shuman et al. (2019) showed that treatments such as BTX-A injections, selective dorsal rhizotomy, and SEMLS minimally affected synergies despite changing the walking patterns. Predictive simulations have the potential to relate synergy complexity to impaired walking abilities, which might help designing specific treatments (e.g., physical therapy protocols) targeting impaired selective motor control.

Bone deformities and resultant altered muscle path trajectories make the use of generic musculoskeletal models linearly-scaled to the subjects’ anthropometry inappropriate for clinical analyses in children with CP. A well established approach to capture these aberrant geometries is through the use of models created from Magnetic Resonance Imaging (MRI) (Arnold et al. (2001); Scheys et al. (2009, 2011a)). Such models have been shown to improve, for example, the accuracy of moment arm estimation in children with CP (Scheys et al. (2011b)). Besides geometries, the muscle-tendon properties are also altered in these children (e.g., smaller muscle volumes and shorter fiber lengths as compared to TD individuals) (Barrett and Lichtwark (2010); Smith et al. (2011); Barber et al. (2011a,b, 2012)). This makes the use of Hill-type muscle-tendon models with generic (i.e., anthropometry-based) parameters unsuited for clinical studies. Indeed, such parameters may not reflect altered muscle force generating capacities and, therefore, result in unrepresentative simulations. To capture the impact of altered muscle-tendon properties on walking performance, the muscle-tendon parameters should be personalized. Different approaches have been proposed for such purpose, including methods based on angle-torque relationships from functional movements (Lloyd and Besier (2003); Falisse et al. (2017)).

Predictive simulations have the potential to shed light upon the influence of altered musculoskeletal properties, impaired selective motor control, and spasticity on walking performance by evaluating the isolated effects of these impairments. Yet only few predictive analyses have used simulations for such purpose. Recent modeling work showed that a musculoskeletal model could reproduce an unimpaired walking pattern with five synergies but not with two synergies similar to those seen after neurological injury, suggesting that impaired control affects walking performance (Meharbi et al. (2019)). Another predictive analysis explored the effects of aging on walking performance by adjusting skeletal and neuromuscular parameters and reported a predominant contribution of loss in muscle strength and mass to reduced energy efficiency (Song and Geyer (2018)). Both studies, however, relied on simple two-dimensional (2D) models, neglecting motor control mechanisms in the frontal plane. To the authors’ knowledge, no study has yet attempted to relate patients’ clinical examination reports to the outcome of predictive simulations evaluating the effects of musculoskeletal and motor control impairments on walking performance based on three-dimensional (3D) subject-specific models.

The purpose of this study was to evaluate the ability of our predictive simulation platform to differentiate the effects of musculoskeletal and motor control impairments on the impaired walking pattern (i.e., crouch gait) of a specific child with CP. To this aim, we evaluated the effect of these impairments on gait patterns predicted by performance optimization (Figure 1A). We first investigated the influence of using personalized rather than generic muscle-tendon parameters, thereby assessing the contribution of the child’s altered muscle-tendon properties to the crouch gait pattern. We then evaluated the impact of imposing a number of synergies lower than typically reported for unimpaired individuals, thereby testing how reducing neuromuscular control complexity affects walking performance. We finally investigated the effect of spasticity modeled based on muscle force and force rate feedback. In all cases, we used a MRI-based musculoskeletal model of the child to take his aberrant geometries into account. We found that the altered muscle-tendon properties rather than the control impairments alone caused a crouch gait pattern. As an additional analysis, we investigated whether the child’s impairments impede a walking pattern similar to TD walking or rather make such a walking pattern less optimal. To this aim, we extended the performance criterion of the predictive simulations with a tracking term that penalized deviations from a TD walking pattern. We found that the musculoskeletal impairments did not prevent an upright walking pattern resembling TD walking but that upright walking was less optimal than walking in crouch.

**Figure 1.**
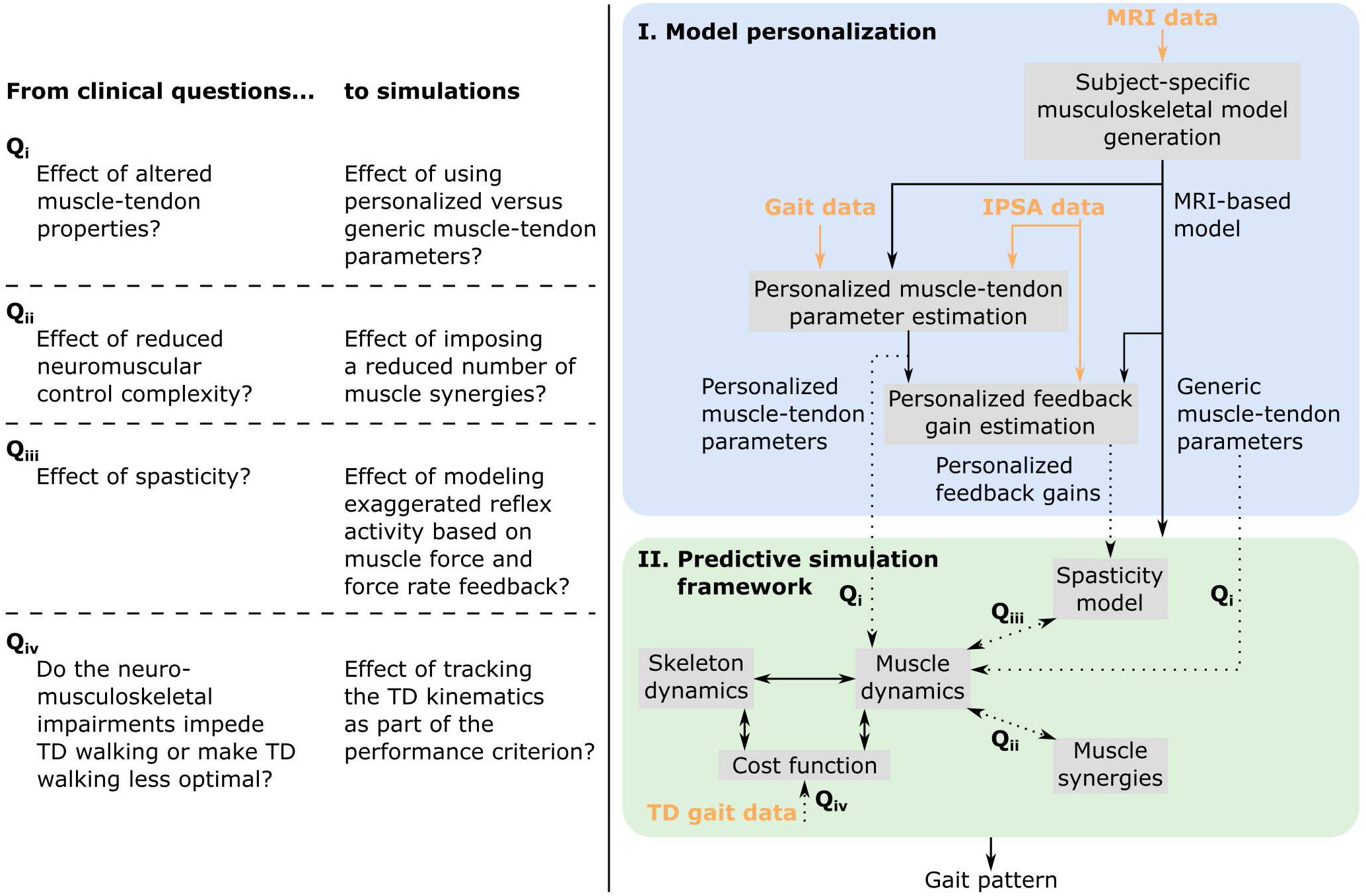
Overview of (A) clinical questions and corresponding simulations, and (B) methodology. MRI images are used to generate a musculoskeletal model of the child with subject-specific geometries. This MRI-based model as well as experimental data collected during walking and instrumented passive spasticity assessments (IPSA) are inputs to optimization procedures providing personalized estimates of Hill-type muscle-tendon parameters characterizing altered muscle-tendon properties and personalized feedback gains characterizing spasticity. The framework for predictive simulations generates gait patterns by optimizing a cost function, describing a walking-related performance criterion, subject to the muscle and skeleton dynamics of the MRI-based musculoskeletal model. We investigated the effects of impairments on predicted gait patterns (dotted arrows): Q_i_ we evaluated the effect of altered versus unaltered muscle-tendon properties by using personalized versus generic muscle-tendon parameters in the muscle dynamics; Q_ii_ we assessed the influence of reducing the neuromuscular control complexity by imposing a reduced number of muscle synergies; Q_iii_ we explored the impact of spasticity on walking performance. Details on how we modeled these impairments are described in the methods. As an additional analysis, Q_iv_, we evaluated how well the model was able to reproduce the gait pattern of a typically developing (TD) child by adding a term in the cost function penalizing deviations between predicted gait pattern and measured gait data of a TD child. All these analyses can be combined as well as performed in isolation. Details are provided in section “model-based analyses”.

## 2 MATERIAL AND METHODS

The overall process to evaluate the effects of impairments on walking performance through predictive simulations is outlined in Figure 1B. The following sections provide details of this process.

### Experimental data

We collected data from one child with diplegic CP (male; age: 15 years; height: 143 cm; mass: 33.1 kg). The data collection was approved by the Ethics Committee at UZ Leuven (Belgium) and written informed consent was obtained from the child’s parents. The child was instrumented with retro-reflective skin mounted markers whose 3D trajectories were recorded (100 Hz) using a motion capture system (Vicon, Oxford, UK) during overground walking at self-selected speed. Ground reaction forces were recorded (1000 Hz) using force plates (AMTI, Watertown, USA). EMG was recorded (2000 Hz) using a telemetric Zerowire system (Cometa, Milan, Italy) from eight muscles of each leg (rectus femoris, biceps femoris short head, semitendinosus, tibialis anterior, gastrocnemius lateralis, vastus lateralis, soleus, and gluteus medius). EMG from the rectus femoris and vastus lateralis was of poor quality and excluded from the analysis.

On the same day as the gait analysis, spasticity of the right medial hamstrings and gastrocnemii was assessed using an instrumented passive spasticity assessment (IPSA; described in detail by Bar-On et al. (2013)). Hamstrings and gastrocnemii were passively stretched by moving knee and ankle, respectively, one at a time from a predefined position throughout the full range of motion (ROM). The stretches were performed at slow and fast velocities. EMG was collected from four muscles (semitendinosus, gastrocnemius lateralis, rectus femoris, and tibialis anterior) using the same system and electrode placement as used for gait analysis. The motion of the distal and proximal segments were tracked using two inertial measurement units (Analog Devices, ADIS16354). The forces applied to the segment were measured using a hand-held six degrees of freedom (DOFs) load-cell (ATI Industrial Motion, mini45). The position of the load-cell relative to the joint axis was manually measured by the examiner.

Muscle strength, selectivity, and ROM were evaluated (Table 1) with a standardized clinical examination protocol (Desloovere et al. (2006)). The child had close to normal ROM at the hip and ankle but bilateral knee extension deficits, bilateral spasticity in most muscles, good strength in most muscles although slight deficits in hip extensors, knee extensors, and hip abductors, and good to perfect selectivity in most muscles. MRI images were collected for the hip region (i.e., pelvis and femur according to the protocol described by Bosmans et al. (2014)). The child was classified at a level II in the Gross Motor Function Classification System (GMFCS).

**Table 1.** Clinical examination. Spasticity, MAS is for Modified Ashworth Scale: 1 is low, 1+ is medium, and 2 is high spastic involvement; Tard is for Tardieu test. Strength: 3 is medium and 4 is good strength; strength from 3 indicates ability to move against gravity. Selectivity: 1 is medium, 1.5 is good, and 2 is perfect selective control. Clinically meaningful deviations from unimpaired individuals are in bold.

|  | Range of motion |  |  | Spasticity |  |
| --- | --- | --- | --- | --- | --- |
|  | Left | Right |  | Left | Right |
| Hip flexion | 145° | 140° | Hip flexion MAS | 2 | 2 |
| Hip extension | -10° | -10° | Hip adduction (Knee 0°) MAS | 1.5 | 1.5 |
| Hip abduction (Knee 0°) | 25° | 25° | Hip adduction (Knee 90°) MAS | 0 | 0 |
| Hip abduction (Knee 90°) | 45° | 45° | Hamstrings MAS | 1.5 | 1 |
| Hip adduction | 0° | 0° | Hamstrings Tard | -70° | / |
| Hip internal rotation (prone) | 60° | 70° | DuncanElly MAS | 1.5 | 1.5 |
| Hip external rotation (prone) | 25° | 25° | DuncanElly Tard | 2° | 2° |
| Hip internal rotation (supine) | 25° | 30° | Soleus MAS | 0 | 0 |
| Hip external rotation (supine) | 55° | 50° | Soleus Tard | / | / |
| Knee flexion | 120° | 120° | Gastrocnemius MAS | 1.5 | 1.5 |
| Knee extension | -20° | -15° | Gastrocnemius Tard | 0° | 5° |
| Knee spontaneous position | -30° | -25° | Tibialis Post MAS | 0 | 0 |
| Popliteal angle unilateral | -70° | -65° | Clonus | 0 | 0 |
| Popliteal angle bilateral | -65° | -60° |  |  |  |
| Ankle dorsiflexion (Knee 90°) | 20° | 25° |  | Alignment |  |
| Ankle dorsiflexion (Knee 0°) | 15° | 15° |  | Left | Right |
| Ankle plantarflexion | 35° | 35° | Femoral anteversion | 35° | 35° |
| Ankle inversion | 40° | 45° | Tibia-femoral angle | 25° | 25° |
| Ankle eversion | 10° | 10° | Bimalleol angle | 40° | 40° |
|  | Selectivity |  |  | Strength |  |
|  | Left | Right |  | Left | Right |
| Hip flexion | 2 | 2 | Hip flexion | 4 | 4 |
| Hip extension | 1.5 | 1.5 | Hip extension | 3 | 3 |
| Hip abduction | 1.5 | 1.5 | Hip abduction | 3+ | 3+ |
| Hip adduction | 2 | 2 | Hip adduction | 4 | 4 |
| Knee flexion | 1.5 | 1.5 | Knee flexion | 4 | 3+ |
| Knee extension | 1 | 1.5 | Knee extension | 3+ | 3+ |
| Ankle dorsiflexion (Knee 90°) | 1.5 | 1.5 | Ankle dorsiflexion (Knee 90°) | 4 | 4 |
| Ankle dorsiflexion (Knee 0°) | 1.5 | 1.5 | Ankle dorsiflexion (Knee 0°) | 4 | 4 |
| Ankle plantarflexion | 1.5 | 1.5 | Ankle plantarflexion | 4 | 3+ |
| Ankle inversion | 1.5 | 1.5 | Ankle inversion | 4 | 4 |
| Ankle eversion | 2 | 1.5 | Ankle eversion | 4 | 4 |

We processed the experimental gait and IPSA data, used as input for the estimation of muscle-tendon parameters and feedback gains (Figure 1; details below), with OpenSim 3.3 (Delp et al. (2007)) using the MRI-based model described below.

### Subject-specific musculoskeletal model generation

A 3D musculoskeletal model with subject-specific geometries was created from MRI images (Scheys et al. (2009, 2011a); Bosmans et al. (2014)). Bones of the lower limbs and pelvis were segmented using Mimics (Materialize, Leuven, Belgium). Anatomical reference frames, joint axes, and muscle origin and insertion points were defined using a previously developed workflow (Scheys et al. (2008)). The model consisted of 21 DOFs (six between the pelvis and the ground; three at each hip joint; one at each knee, ankle, and subtalar joint; and three at the lumbar joint), 86 muscles actuating the lower limbs (43 per leg), three ideal torque actuators at the lumbar joint, and four contact spheres per foot (Delp et al. (1990, 2007)). We added passive torques to the joints of the lower limbs and the trunk to model the role of the ligaments and other passive structures (Anderson and Pandy (2001)). These passive torques varied exponentially with joint positions and linearly with joint velocities.

We used Raasch’s model (Raasch et al. (1997); De Groote et al. (2009)) to describe muscle excitation-activation coupling (muscle activation dynamics) and a Hill-type muscle-tendon model (Zajac (1989); De Groote et al. (2016)) to describe muscle-tendon interaction and the dependence of muscle force on fiber length and velocity (muscle contraction dynamics). We modeled skeletal motion with Newtonian rigid body dynamics and smooth approximations of compliant Hunt-Crossley foot-ground contacts (Delp et al. (2007); Sherman et al. (2011); Falisse et al. (2019)). We calibrated the Hunt-Crossley contact parameters (transverse plane locations and contact sphere radii) through muscle-driven tracking simulations of the child’s experimental walking data as described in previous work (Falisse et al. (2019)). To increase computational speed, we defined muscle-tendon lengths, velocities, and moment arms as a polynomial function of joint positions and velocities (van den Bogert et al. (2013); Falisse et al. (2019)).

### Personalized muscle-tendon parameter estimation

The force-length-velocity relationships describing the force generating capacity of the Hill-type muscle-tendon model are dimensionless and can be scaled to a specific muscle through five muscle-tendon parameters: the maximal isometric force 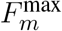, the optimal fiber length 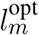, the tendon slack length 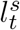, the optimal pennation angle 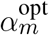, and the maximal fiber contraction velocity 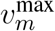 (assigned to ten times 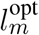). In this study, we used generic and personalized parameters when generating predictive simulations of walking (Figure 1).

The generic parameters were derived by linearly scaling the parameters of a generic musculoskeletal model (Delp et al. (1990)) to the child’s anthropometry. The linear scaling was only performed for the optimal fiber lengths and tendon slack lengths. The maximal isometric muscle forces were scaled based on body mass *M* (van der Krogt et al. (2016)):

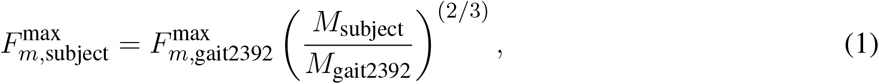

where gait2392 refers to the OpenSim gait2392 model (Delp et al. (1990, 2007)).

The personalized parameters reflect the muscle force generating capacity of the subject. Only optimal fiber lengths and tendon slack lengths were personalized as gait simulations have been shown to be the most sensitive to these two parameters (De Groote et al. (2010)). The personalization process was based on an extension of an optimal control approach to solve the muscle redundancy problem while accounting for muscle dynamics (De Groote et al. (2016); Falisse et al. (2017)). Solving the muscle redundancy problem identifies muscle excitations that reproduce joint torques underlying a given movement while minimizing a performance criterion (e.g., muscle effort). We augmented this formulation in different ways. First, we added optimal fiber lengths and tendon slack lengths as optimization variables. Second, we introduced a term in the cost function minimizing the difference between muscle activations and scaled EMG signals where scale factors were included as optimization variables. Third, we assumed that muscles operate around their optimal fiber lengths, and that maximal and minimal fiber lengths across movements should hence be larger and smaller, respectively, than their optimal fiber lengths. Fourth, we assumed that resistance encountered when evaluating the ROM during the clinical examination may be, at least in part, attributed to passive muscle forces. Hence, we included a term in the cost function minimizing the difference between fiber lengths at these extreme positions of the ROM and reference fiber lengths generating large passive forces. Finally, we minimized optimal fiber lengths, assuming that children with CP have short fibers (Barrett and Lichtwark (2010)). The problem thus consisted in identifying muscle excitations and parameters that minimized a multi-objective cost function:

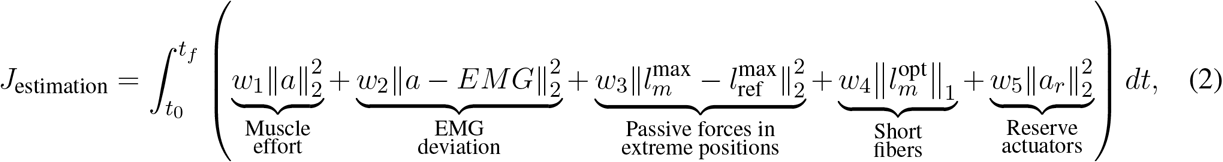

where *t*_0_ and *t*_*f*_ are initial and final times, *a* are muscle activations, 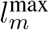 and 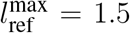 are simulated and reference fiber lengths, respectively, at the extreme positions of the ROM, *a*_*r*_ are reserve actuators, *w*_1−5_ are weight factors, and *t* is time. This cost function was subject to constraints enforcing muscle dynamics, that resultant muscle forces should reproduce joint torques calculated from inverse dynamics, that fiber lengths should cross their optimal fiber lengths during the movement, and that the difference between activations and EMG should not be larger than 0.1. Reserve actuators are non-physiological ideal actuators added to muscle-generated torques to ensure that joint torques from inverse dynamics can be reproduced. The weights were manually adjusted to the following: *w*_1_ = 10 × 10^−4^, *w*_2_ = 30 × 10^−4^, *w*_3_ = 3550 × 10^−4^, *w*_4_ = 1010 × 10^−4^, and *w*_5_ = 5400 × 10^−4^. These weights primarily penalized the use of reserve actuators and encouraged the generation of passive forces in the extreme positions of the ROM. We solved this problem while simultaneously considering data from four gait trials of each leg and six passive stretches (IPSA measurements) of the right hamstrings, rectus femoris, and gastrocnemii at slow and fast velocities (one stretch per muscle per speed). Data from 14 trials (gait and passive trials combined) was thus included. Data from passive stretches of left leg muscles was not available. Hence, we imposed that corresponding parameters of both legs could not differ by more than 5%. The parameters were allowed to vary between 50 and 200% of the generic values.

### Spasticity model - Personalized feedback gain estimation

We modeled spasticity through delayed feedback from muscle-tendon force and its first time derivative (i.e., force rate) (Falisse et al. (2018)). The model relates sensory information *s* (i.e., muscle force and force rate) to feedback muscle activations *a*_*s*_ through a first order differential equation:

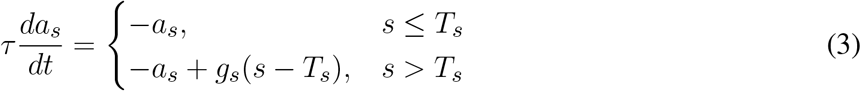

where *T*_*s*_ is a feedback threshold, *g*_*s*_ is a feedback gain, and *τ*_*s*_ = 30 ms is a time delay.

We determined the threshold for force feedback as the value 20 ms before the EMG onset (Staude and Wolf (1999)) and used a zero threshold for force rate feedback. We identified the personalized feedback gains that minimized the difference between EMG and feedback muscle activations during fast passive stretches (IPSA measurements). We performed such optimization for the right medial hamstrings (i.e., biceps femoris long head, semitendinosus, and semimembranosus) and for the right gastrocnemii (i.e., gastrocnemius lateralis and medialis). We used semitendinosus EMG to drive the three hamstrings and gastrocnemius lateralis EMG to drive both gastrocnemii. We normalized EMG using scale factors identified when estimating the personalized muscle-tendon parameters. We described the optimization process in detail in previous work (Falisse et al. (2018)). We incorporated the spasticity model with personalized feedback gains in our framework for predictive simulations (Figure 1). Since we only had IPSA measurement for the right leg, we used feedback gains and thresholds identified with right leg data for left leg muscles. Gait EMG data and spasticity, as clinically assessed (Table 1), were comparable for both legs.

### Muscle synergies

We modeled the reduced neuromuscular control complexity through muscle synergies. These synergies consisted of two matrices: a *N*_syn_ × *N*_*f*_ matrix *H*, where *N*_syn_ is the number of synergies and *N*_*f*_ is the number of frames, containing synergy activations and a *N*_*m*_ × *N*_syn_ matrix *W*, where *N*_*m*_ is the number of muscles, containing weights that determine the contribution of each muscle in each synergy. Individual muscle activations were composed from synergies as follows:

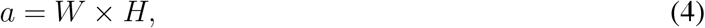

where *a* has dimensions *N*_*m*_ × *N*_*f*_. Importantly, we did not impose subject-specific synergies when generating predictive simulations (Figure 1). Instead, we modeled the effect of reducing the neuromuscular control complexity by limiting the number of synergies per leg to four or three, thereby limiting the selection of independent muscle activations. This represents a reduction of the neuromuscular control complexity under the assumption that five synergies describe healthy human locomotion (Ivanenko et al. (2004)).

### Problem formulation

We predicted gait patterns by optimizing a gait-related cost function, independent of experimental data, based on the MRI-based musculoskeletal model described above. In addition to optimizing performance, we imposed average gait speed and periodicity of the gait pattern. We optimized for a full gait cycle to account for asymmetry of CP gait. We solved the resultant optimal control problem via direct collocation. The problem formulation and computational choices are detailed in previous work (Falisse et al. (2019)).

The cost function represents the goal of the motor task. We modeled this task-level goal as a weighted sum of gait-related performance criteria including metabolic energy rate, muscle fatigue, joint accelerations, passive joint torques, and trunk actuator excitations:

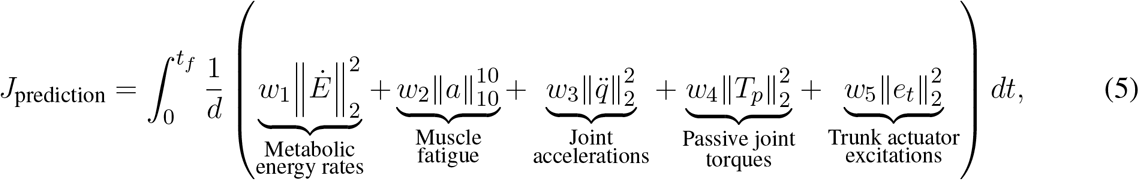

where *t*_*f*_ is unknown gait cycle duration, *d* is distance travelled by the pelvis in the forward direction, 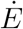 are metabolic energy rates, *a* are muscle activations, 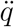 are joint accelerations, *T*_*p*_ are passive joint torques, *e*_*t*_ are excitations of the trunk torque actuators, *w*_1−5_ are weight factors, and *t* is time. We modeled metabolic energy rate using a smooth approximation of the phenomenological model described by Bhargava et al. (2004). This metabolic model requires parameters for fiber type composition and muscle specific tension, which we obtained from the literature (Uchida et al. (2016)). We manually adjusted the weight factors until we found a set of weights that predicted human-like walking: *w*_1_ = (25/86/body mass) × 10^−2^, *w*_2_ = 25/86 × 10^2^, *w*_3_ = 50/21, *w*_4_ = 10/15 × 10^2^, and *w*_5_ = 1/3 × 10^−1^. We added several path constraints enforcing a prescribed average gait speed corresponding to the child’s average gait speed (*d/t*_*f*_ = 1 m s^−1^), imposing periodic states over the complete gait cycle (except for the pelvis forward position), and preventing inter-penetration of body segments.

### Model-based analyses

We investigated the differential effects of altered muscle-tendon properties, reduced neuromuscular control complexity, and spasticity on gait patterns predicted with the MRI-based musculoskeletal model (Figure 1). In particular, we compared predicted joint kinematics and kinetics, muscle activity, and stride lengths to their experimental counterparts. We also evaluated how impairments affected the metabolic cost of transport (COT), defined as metabolic energy consumed per unit distance traveled.

First, we tested the influence of altered versus unaltered muscle-tendon properties by using personalized versus generic muscle-tendon parameters in the muscle dynamics (Q_i_ in Figure 1). In this initial analysis, we did not include spasticity, nor imposed synergies.

Second, we assessed the impact of reducing the neuromuscular control complexity by imposing fixed numbers of synergies (Q_ii_ in Figure 1). To assess the effect of reducing the number of synergies, we compared the synergy activations resulting from simulations with three and four synergies using the coefficient of determination R^2^ and the synergy weights using Pearson’s coefficient of correlation *r*. We generated simulations with both sets of muscle-tendon parameters to explore the effect of synergies in isolation as well as in combination with altered muscle-tendon properties.

Finally, we evaluated the effect of spasticity in the three medial hamstrings and two gastrocnemii of both legs (Q_iii_ in Figure 1). We modeled muscle activations as the sum of reflex muscle activations determined based on the personalized spasticity model and feedforward muscle activations:

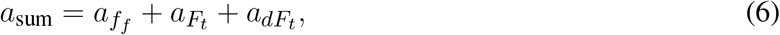

where 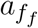 are feedforward muscle activations, and 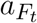 and 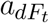 are muscle activations from muscle force and force rate feedback, respectively. We only tested the effect of spasticity based on the model with personalized muscle-tendon parameters, since these parameters were used to estimate the feedback gains. We tested the effect of spasticity in combination with selective control (i.e., no synergy constraints) as well as with a reduced number of muscle synergies.

As an additional analysis, we investigated whether the child adopted an impaired crouch gait pattern because of neuro-mechanical constraints or because it was more optimal (Q_iv_ in Figure 1). To this aim, we added a term in the cost function that penalized deviations from measured kinematics of a TD child:

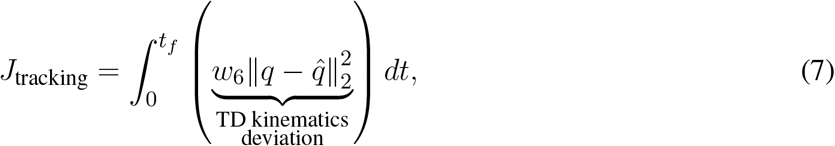

where 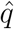 are measured joint positions of a TD child and *w*_6_ = 100/20 is a weight factor. We generated these simulations with personalized parameters as well as with and without synergies. We did not include spasticity in this analysis since it had little influence on the walking pattern in the simulations described above.

We formulated our problems in MATLAB using CasADi (Andersson et al. (2019)), applied direct collocation using a third order Radau quadrature collocation scheme with 150 mesh intervals per gait cycle, and solved the resulting nonlinear programming problems with the solver IPOPT (Wächter and Biegler (2006)). We applied algorithmic differentiation to compute derivatives (Falisse et al. (2019)). We started each optimization from multiple initial guesses and selected the result with the lowest optimal cost. Initial guesses for joint variables were based on experimental data. Specifically, for all simulations, we used two initial guesses derived from experimental kinematics of the CP and the TD child. For simulations accounting for synergies, we added initial guesses derived from simulated kinematics with the lowest optimal costs produced without synergies and with more synergies (e.g., with three synergies, initial guesses were derived from the best kinematic solutions with four synergies and without synergies). For simulations accounting for spasticity, we added initial guesses derived from simulated kinematics with the lowest optimal costs produced without spasticity. In all cases, initial guesses for muscle, trunk, and synergy variables were constant across time and not informed by experimental data. Initial guesses for synergy weights were constant across muscles and independent of experimental data.

## RESULTS

### Gait analysis

The child walked with a pronounced crouch gait pattern characterized by bilateral knee extension deficits with reduced knee ROM during swing, a lack of right ankle dorsiflexion at the end of swing, excessive left ankle dorsiflexion, excessive and deficient right and left hip adduction, respectively, and excessive bilateral hip internal rotation (Figures 2 and S1).

**Figure 2.**
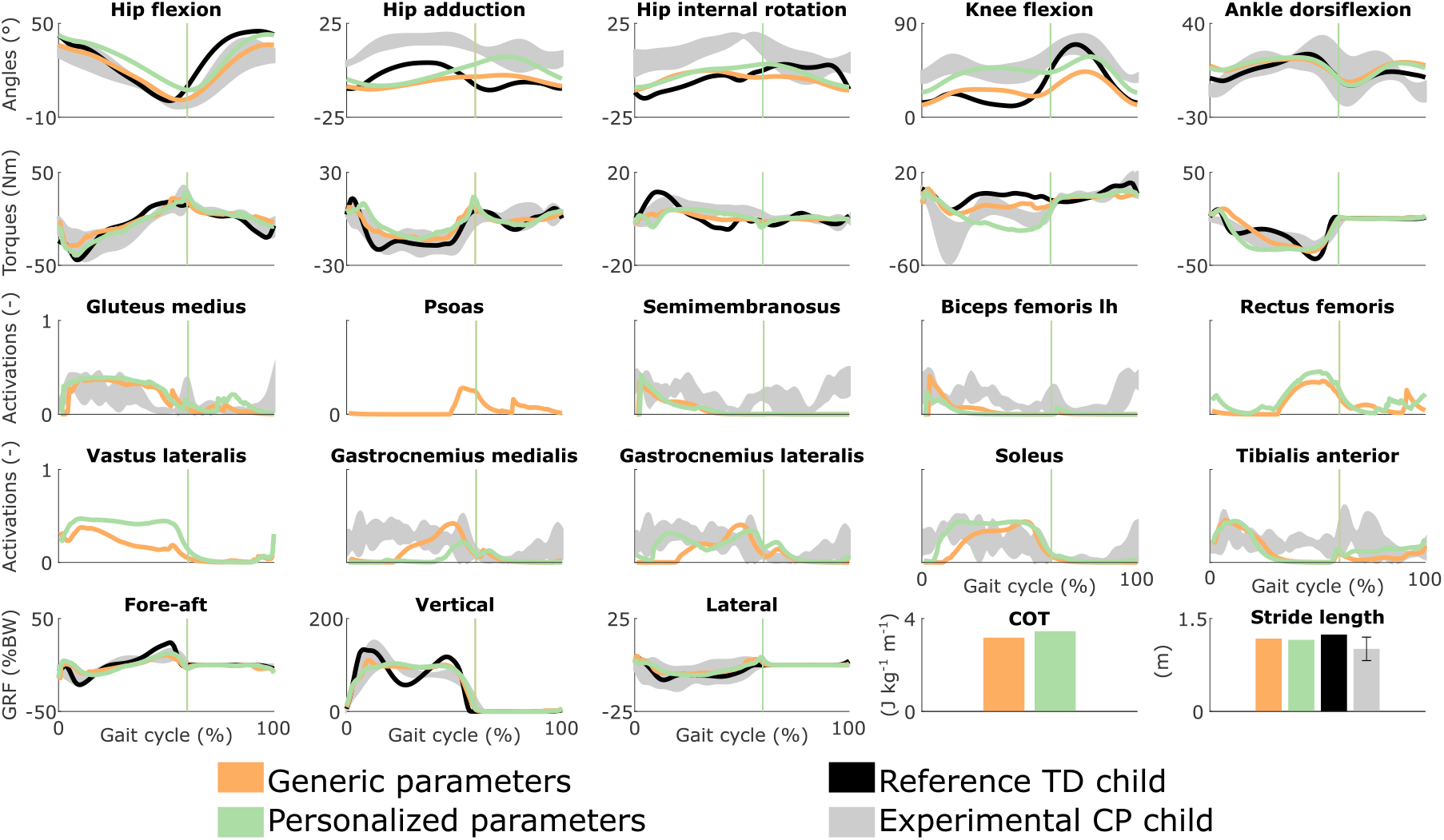
Influence of the muscle-tendon parameters on the predicted walking gaits. Variables from the right leg are shown over a complete gait cycle; left leg variables are shown in Figure S1 (Supplementary Material). Vertical lines indicate the transition from stance to swing. Experimental data is shown as mean two standard deviations. Experimental EMG data was normalized to peak activations. GRF is for ground reaction forces; BW is for body weight; COT is for metabolic cost of transport; lh is for long head.

### Influence of the muscle-tendon parameters

Using personalized versus generic muscle-tendon parameters resulted in a crouch (i.e., excessive knee flexion) versus a more upright gait pattern (Figures 2 and S1; Movies S1-2). Personalized optimal fiber lengths and tendon slack lengths were generally smaller and larger, respectively, than their generic counterparts (Tables S1-2). The use of personalized parameters resulted in decreased deviations (smaller root mean square error (RMSE)) between measured and predicted knee angles (RMSE of 17° and 11° for the left and right leg, respectively) as compared to the use of generic parameters (RMSE of 43° and 25°). The gastrocnemius lateralis and soleus (ankle plantarflexors) were activated earlier in stance with the crouch gait, as observed in the child’s EMG. The vasti (knee extensors) activity was also increased during stance when the model walked in crouch. The COT was higher with the personalized parameters (crouch gait; 3.45 J kg^−1^m^−1^) than with the generic parameters (more upright gait; 3.18 J kg^−1^m^−1^). Predicted stride lengths were larger than the average stride length of the child but were within two standard deviations.

### Influence of the synergies with generic muscle-tendon parameters

Reducing the number of synergies in combination with generic muscle-tendon parameters did not induce the amount of crouch that was experimentally measured in the child, although it altered muscle coordination and increased COT (Figures 3 and S2; Movie S3). The right knee flexion angles increased during stance with the reduction of the neuromuscular control complexity but were still smaller than experimentally measured. This was accompanied with increased rectus femoris (knee extensor) activity. The synergies had a limited effect on the left leg that had a straight knee pattern during stance. The COT increased with the reduction of the neuromuscular control complexity (3.58 and 3.90 J kg^−1^m^−1^ with four and three synergies, respectively). The synergies had little effect on the predicted stride lengths that were larger than the child’s average stride length but were within two standard deviations. The synergies of the three-synergy case were similar to the first three synergies of the four-synergy case (average R^2^ and *r* over three common synergy activations and weight vectors, respectively, of both legs: 0.84 ± 0.19 and 0.83 ± 0.10). The additional synergy in the four-synergy case was activated in early stance and at the transition between stance and swing, and mainly consisted of hip adductors.

**Figure 3.**
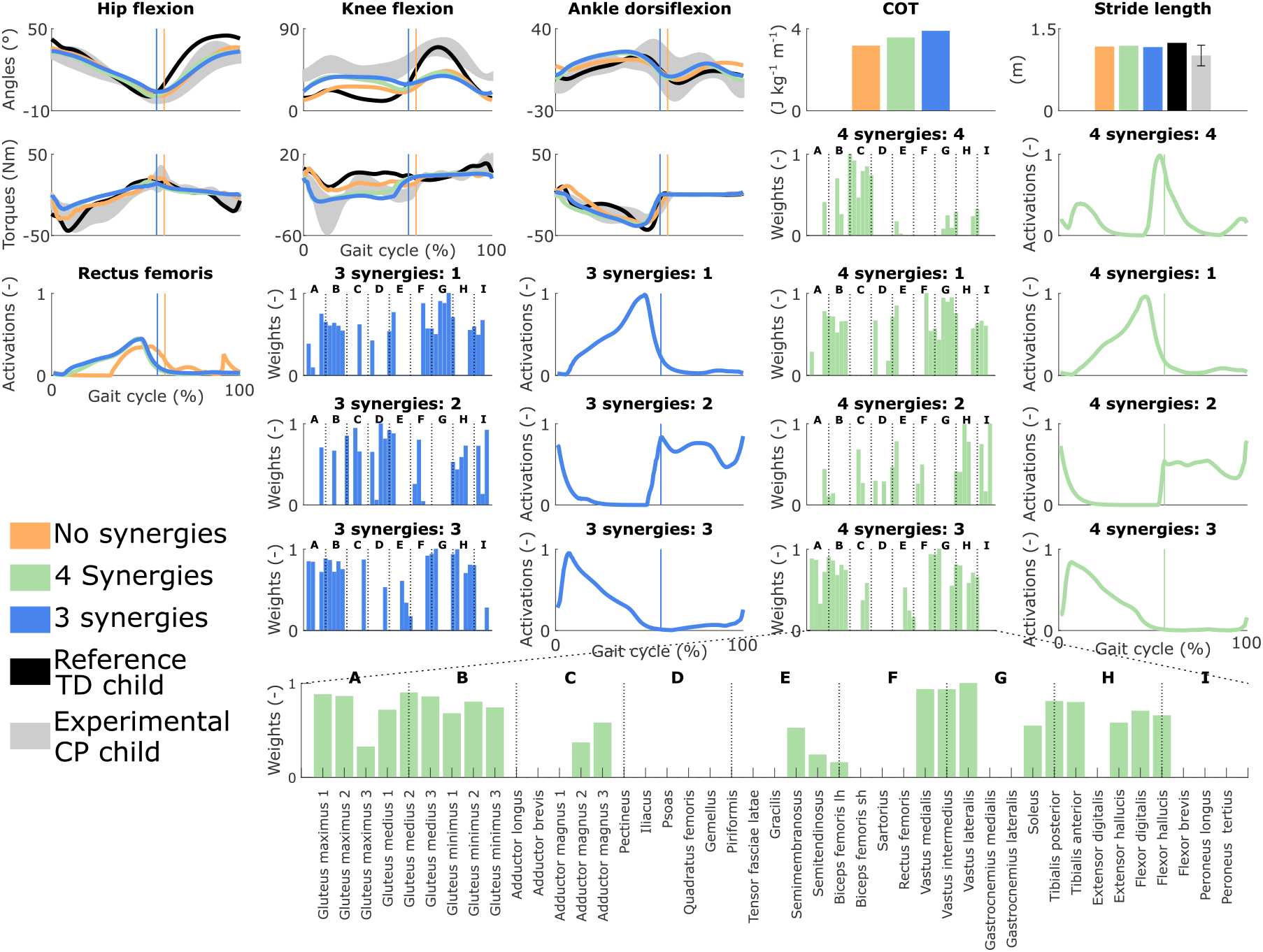
Influence of the synergies on walking gaits predicted with the generic muscle-tendon parameters. Variables from the right leg are shown over a complete gait cycle; left leg variables are shown in Figure S2 (Supplementary Material). Vertical lines (solid) indicate the transition from stance to swing. Panels of synergy weights are divided into sections (A-I) to relate bars to muscle names provided in the bottom bar plot, which is an expanded version of the plot of weights with title 4 synergies: 3. Lh and sh are for long and short head, respectively. Weights were normalized to one. Experimental data is shown as mean ± two standard deviations.

### Influence of the synergies with personalized muscle-tendon parameters

Reducing the number of synergies in combination with personalized muscle-tendon parameters had a minor effect on gait kinematics but altered muscle coordination and increased COT (Figures 4 and S3; Movie S4). Specifically, synergies only had a slight effect on the kinematics during the swing phase of the right leg but affected the activation pattern of certain muscles (e.g., gastrocnemius medialis and lateralis). The COT increased with the reduction of the neuromuscular control complexity (3.94 and 4.09 J kg^−1^m^−1^ with four and three synergies, respectively). Stride lengths slightly decreased with synergies but remained larger than the child’s average stride length. The synergies of the three-synergy case were similar to the first three synergies of the four-synergy case (average R^2^ and *r*: 0.85 ± 0.05 and 0.87 ± 0.09, respectively). The additional synergy in the four-synergy case was activated in early stance and at the transition between stance and swing, and mainly consisted of the gemellus, piriformis, tibialis posterior, and several ankle plantarflexors.

**Figure 4.**
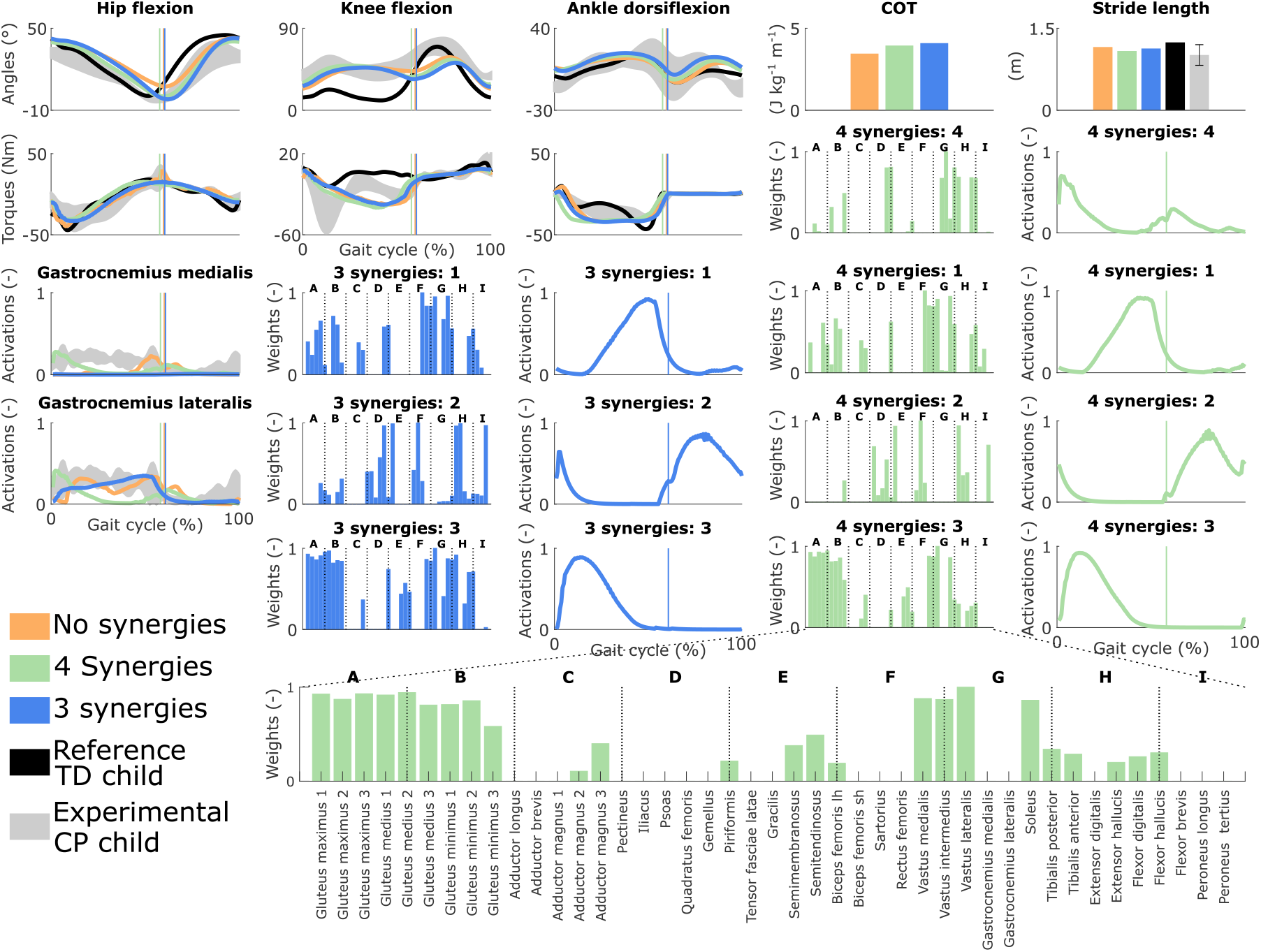
Influence of the synergies on walking gaits predicted with the personalized muscle-tendon parameters. Variables from the right leg are shown over a complete gait cycle; left leg variables are shown in Figure S3 (Supplementary Material). Vertical lines (solid) indicate the transition from stance to swing. Panels of synergy weights are divided into sections (A-I) to relate bars to muscle names provided in the bottom bar plot, which is an expanded version of the plot of weights with title 4 synergies: 3. Lh and sh are for long and short head, respectively. Weights were normalized to one. Experimental data is shown as mean ± two standard deviations. Experimental EMG data was normalized to peak activations.

### Influence of spasticity

Spasticity had a limited effect on muscle coordination and almost no influence on gait kinematics (Figures 5 and S4; Movie S5). Specifically, spastic activity was predicted in the medial hamstrings in early stance but this had, overall, a minor effect on the total (i.e., combined spastic and non-spastic contributions) medial hamstrings activity when compared to simulations without spasticity. Bursts of spastic activity were also observed in early swing. Medial hamstrings activity contributes to knee flexion but since similar (timing and magnitude) activity profiles were predicted with and without spasticity, there was no difference in predicted knee flexion angles. A constant low spastic contribution was predicted for the gastrocnemius lateralis during stance, whereas a minor contribution was predicted for the gastrocnemius medialis during stance and at the transition between stance and swing. Spasticity hence does not explain the lack of right ankle dorsiflexion (i.e., increased plantarflexion) observed at the end of swing in experimental data. Similar observations hold with and without synergies. The COT increased when incorporating spasticity (3.75 and 4.18 J kg^−1^m^−1^ with zero and four synergies, respectively).

**Figure 5.**
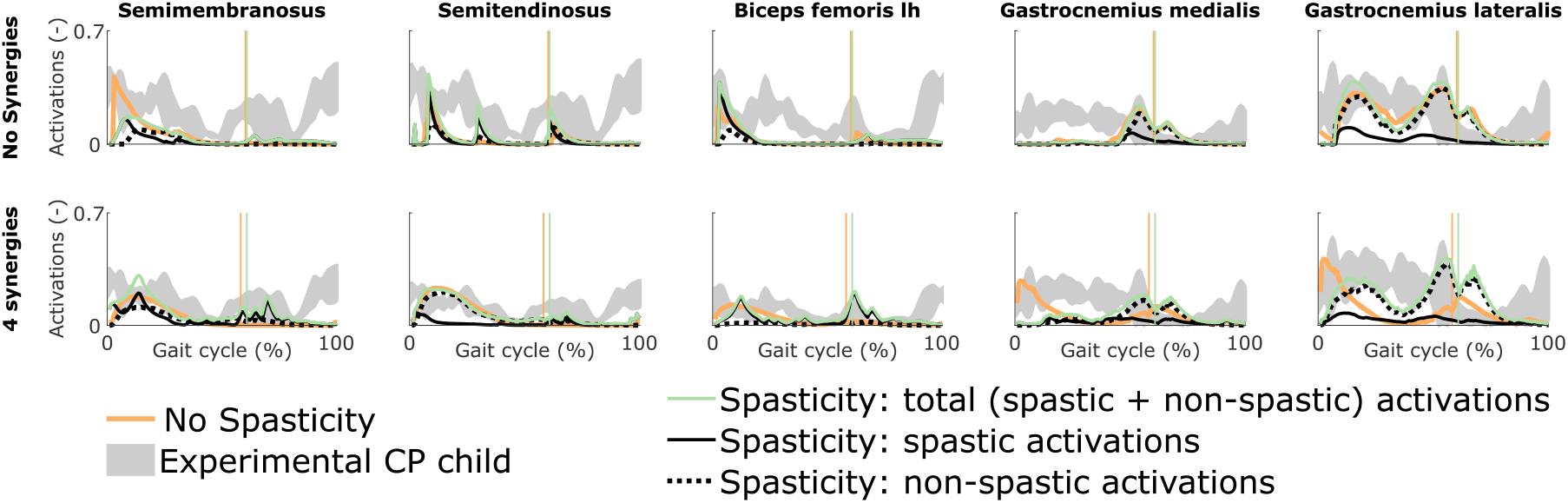
Influence of spasticity on the predicted muscle activity. Activations from right leg muscles only are shown over a complete gait cycle; left leg activations are shown in Figure S4 (Supplementary Material). When accounting for spasticity, total activations (green) combine spastic (solid black) and non-spastic (dotted black) activations. Vertical lines indicate the transition from stance to swing. Experimental data is shown as mean ± two standard deviations. Experimental EMG data was normalized to peak activations. Lh is for long head.

### Influence of tracking the kinematics of a TD child

Tracking the TD kinematics while using personalized muscle-tendon parameters produced an upright gait pattern when not incorporating synergies, but decreased the overall gait performance (Figures 6 and S5; Movie S6). Specifically, the simulated gait had a similar COT (3.46 J kg^−1^m^−1^) as the crouch gait pattern predicted without such tracking term but the contribution of most terms in the cost function increased, suggesting that walking upright is not prevented by mechanical constraints (i.e., aberrant musculoskeletal geometries and altered muscle-tendon properties) but is less optimal, due to these mechanical constraints, than walking in crouch for this child. The contribution of the muscle fatigue term increased by 29%, in part driven by higher activations of the glutei. The contribution of the joint acceleration, metabolic energy rate, and passive joint torque terms increased by 15, 15, and 36%, respectively, when walking upright. Similarly, passive muscle forces increased when walking upright for the iliacus and psoas (hip flexors), and biceps femoris short head (knee flexor). Knee flexion increased when adding synergies but did not reach the angle that was experimentally measured in the child (Figure S6). Nevertheless, this suggests that reduced neuromuscular control complexity may contribute to crouch gait. The gastrocnemius lateralis and soleus (ankle plantarflexors) were also activated earlier during stance with synergies. Imposing synergies increased the COT (4.12 and 4.05 J kg^−1^m^−1^ with four and three synergies, respectively).

**Figure 6.**
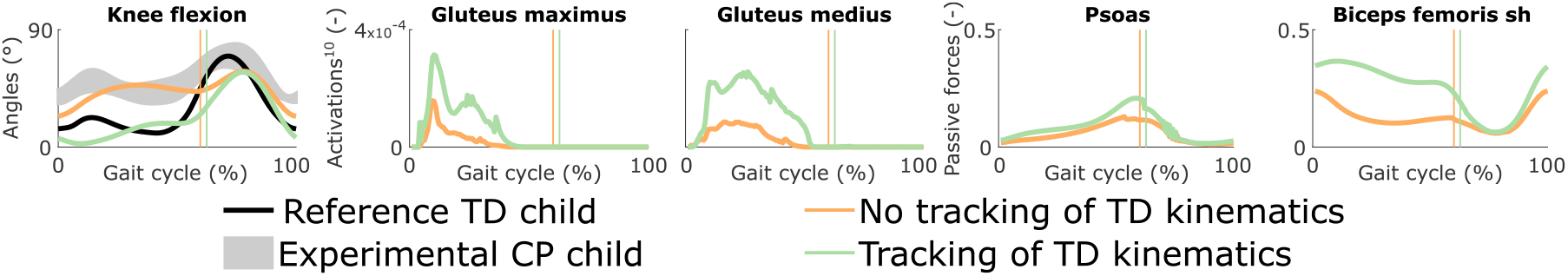
Influence of tracking the TD kinematics on predicted walking gaits. Variables from the right leg are shown over a complete gait cycle; left leg variables are shown in Figure S5 (Supplementary Material). Vertical lines indicate the transition from stance to swing. Experimental data is shown as mean ± two standard deviations. Muscle fatigue is modeled by activations at the tenth power. Passive muscle forces are normalized by maximal isometric muscle forces. Sh is for short head. The influence of synergies on predicted walking gaits is depicted in Figure S6 (Supplementary Material).

## DISCUSSION

We demonstrated the ability of predictive simulations to explore the differential effects of musculoskeletal and motor control impairments on the gait pattern of a child with CP. In this specific case, aberrant musculoskeletal geometries and altered muscle-tendon properties explained the key gait deviation of the child, namely the crouch gait pattern. Accounting for aberrant geometries alone (i.e., MRI-based model with generic muscle-tendon parameters) did not result in a crouch gait pattern. Despite altered muscle-tendon properties and aberrant geometries, the model could still adopt a more upright gait pattern (TD kinematics tracking). Yet such pattern was less optimal as it induced higher muscle fatigue compared to the crouch gait pattern. These simulations thus suggest that adopting an upright gait pattern for this child might produce an early onset of fatigue, which might explain in part why the child walks in crouch. Importantly, not only fatigue, but also joint accelerations, passive joint torques, and metabolic energy rates increased with an upright gait pattern, likely contributing to the selection of a crouch gait pattern.

Decreasing the neuromuscular control complexity through a reduced number of synergies had a lower effect on the simulated gait patterns than muscular deficits as evaluated when comparing simulated gait patterns obtained with personalized and generic muscle-tendon parameters. Nevertheless, the synergies resulted in increased knee flexion in several simulations, indicating that impaired selective motor control may contribute to gait deficits as suggested in prior simulation studies (Meharbi et al. (2019)). In this study, we imposed the number of synergies but not the synergy structure (synergy weights and activations were optimization variables and not informed by experimental data). We thus explored the effect of reducing the neuromuscular control complexity but not the impact of imposing the child’s experimental synergies. We expect this impact to be limited for this child since he had a good selectivity. Nevertheless, further work should consider such investigation.

Our predictive simulations generated both movement patterns and the underlying synergies. Only imposing the number of synergies resulted in synergies that presented common features with those reported in the literature, such as one synergy activated during early stance and composed by the glutei and vasti, and one synergy activated during late stance consisting of the glutei, ankle plantarflexors, and iliacus (De Groote et al. (2014)). This suggests that synergy structures might emerge from mechanical constraints and performance optimization during walking. Future research should explore this hypothesis based on a larger population.

Decreasing the number of synergies resulted in a larger COT, as may be expected with a higher level of co-activations. This finding has been hypothesized in previous studies (Steele et al. (2017); Meharbi et al. (2019)) but not tested explicitly. It is indeed difficult to dissociate the influence of the neuromuscular control complexity on the COT through experiments or based on measured data, since many other factors (e.g., spasticity (Hemingway et al. (2001)) and weakness (van der Krogt et al. (2012))) might also play a role. Overall, our predictive simulations allow exploring the effects of isolated impairments on gait energetics, which was not possible through analyses based on measured data.

Spasticity had a minor influence on the predicted gait kinematics, suggesting a low impact of spasticity on gait performance for this child. This hypothesis is in agreement with severeal studies reporting a lack of correlation between spasticity as diagnosed during passive movements and determinants of gait (Ada et al. (1998); Marsden et al. (2012); Willerslev-Olsen et al. (2014)). However, it would be premature to draw such conclusion based on this analysis for a single child. First, spasticity was only taken into account for the medial hamstrings and gastrocnemii, whereas the rectus femoris and several hip flexors and adductors were also reported to be spastic (Table 1). Including these other muscles may have an influence on walking performance. Second, experimental data from the spasticity assessment was only collected for the right leg, whereas bilateral spasticity was reported (Table 1). We optimized the feedback parameters using that data but used the resulting parameters for both legs, which might affect our predictions. Third, we used feedback parameters optimized from passive stretches to predict spasticity (i.e., reflex activity) during gait, assuming no reflex modulation. This assumption is in line with the decreased reflex modulation reported for patients with spasticity (Sinkjaer et al. (1996); Faist et al. (1999); Dietz (2002); Dietz and Sinkjaer (2007)). Yet further research is needed to ensure that the same model is valid in passive and active conditions. Finally, the optimized feedback gains depend on EMG that was normalized using scale factors optimized during the muscle-tendon parameter estimation. However, these factors may not truly reflect the magnitude of the spastic responses, which may result in an under- or over-estimation of the predicted spastic activity during gait. In previous work (Falisse et al. (2018)), we showed that predicted spastic responses of the gastrocnemii were in agreement with large EMG signals observed in early stance in subjects with an equinus gait (i.e., toe walking). Interestingly, in this study, the child walked on his toes but we did not observe such EMG rise. Hence, our model predictions were in agreement with the lack of gastrocnemius EMG activity observed during early stance.

Our analysis suggests that muscle-tendon properties rather than selective motor control and spasticity should be the target of interventions aiming to restore an upright posture for this child. This suggestion is in line with the surgical report and one-year post-operative gait analysis. Specifically, the child underwent SEMLS consisting of bilateral rectus femoris transfer, distal femur extension and derotation osteotomy, tibia derotation, and patella distalization that successfully addressed the knee extension deficits and restored the upright gait pattern. The intervention also included bilateral BTX-A injections in the psoas (hip flexor) and gracilis (hip flexor, adductor, and knee flexor) to reduce spasticity. However, BTX-A injections are unlikely to have had an effect one year post-treatment (Molenaers et al. (2010)), suggesting a limited contribution of reduced psoas and gracilis spasticity on restored knee extension. Note that our study did not investigate the sensitivity of the predicted walking patterns to bone misalignment as we considered the same aberrant geometries for all analyses. Studying the effect of bone deformities on the gait pattern should be considered in future work.

Our simulations with personalized muscle-tendon parameters captured salient features of the child’s walking pattern. Nevertheless, they deviated from measured data in different ways. In particular, our model did not adopt the observed equinus gait. Such pattern might have different underlying roots. On the one hand, it might be an ankle strategy to add functional limb length and compensate for the knee extension deficits. Our simulations did not predict such compensation strategy but also lacked knee flexion in early stance as compared to measured data (Figure 2). Increased knee flexion might strengthen the need for ankle compensation, causing the model to adopt an equinus gait. On the other hand, it might be due to contracture of the plantarflexors (Wren et al. (2005); Mathewson et al. (2015)) although this hypothesis is less likely for this child who had a normal ROM in terms of plantarflexion.

Other factors might have contributed to the deviations between predicted and measured movements. First, the musculoskeletal model had generic rather than subject-specific (i.e., MRI-based) geometries for feet and tibias. Since the child later underwent a surgery that included bilateral tibia derotation, these generic geometries might have contributed to the gait deviations. Second, the clinical examination indicated that the child’s trunk was leaning forward. This is likely a compensation strategy, since no fixed lordosis was reported. However, our model had a very simple trunk representation (i.e., one joint with three DOFs), limiting the emergence of compensation strategies. Hence, our simulations resulted in an upright trunk posture, whereas a forward leaning posture might have caused an equinus gait. How to model the trunk to capture such compensations remains an open question. Third, our control strategy likely did not capture all complex control mechanisms that might be at play during gait. For instance, we did not consider in our cost function criteria such as head stability (Menz et al. (2003)) and pain that might contribute to gait control. Further, we designed our cost function based on previous work with a healthy adult but the same performance criterion might not hold for children with CP. Nevertheless, our cost function predicted, as expected, a crouch gait pattern with personalized parameters and a more upright gait pattern with generic parameters, suggesting that it captured at least part of the child’s control strategy. Finally, the personalized muscle-tendon parameters might not accurately capture the effect of the child’s altered muscle-tendon properties. In previous work (Falisse et al. (2017)), we underlined the importance of incorporating experimental data from multiple functional movements when calibrating muscle-tendon parameters in order to obtain valid parameter estimates (i.e., representative of the subject). In this study, the available experimental data was limited to walking trials and passive stretches from one leg. Hence, it is likely that some parameters were calibrated to fit the experimental data but did not truly reflect the force-generating capacities of the child. When used in conditions different from the experiments, these parameters may hence result in non-representative force predictions. A challenge for upcoming research will be the design of experimental protocols to collect experimental data that contains sufficient information for providing valid muscle-tendon parameter estimates while accounting for physiological limitations of impaired individuals and practical limitations of clinical contexts. It is also worth noting that our parameter estimation procedure only adjusted optimal fiber lengths and tendon slack lengths, whereas other parameters may need to be personalized, such as maximal isometric muscle forces, tendon compliance, or maximal muscle contraction velocities. The muscle force-length-velocity relationships might also be altered in children with CP due to their longer sarcomere lengths. Overall, further tuning of the neuro-musculoskeletal model and validation of the simulation framework outcome with a large population are necessary for augmenting the representativeness of the simulations.

## CONCLUSION

This study suggests that predictive simulations are able to identify the main treatment targets for specific patients. In particular, our results showed that, in the presence of aberrant musculoskeletal geometries, altered muscle-tendon properties rather than reduced neuromuscular control complexity and spasticity were the primary driver of the impaired crouch gait pattern observed for the child with CP of this study. Based on this observation, we would recommend altered muscle-tendon properties to be the primary target of clinical interventions aiming to restore a more upright posture, which is in line with the surgical report and one-year post-operative gait analysis. Validation of our simulation workflow through analysis of many more cases is, however, necessary to build confidence in the simulation outcomes. Overall, these results open the door for predicting the functional outcome of treatments on walking performance by allowing *in silico* assessment of the effect of changes in the neuro-musculoskeletal system on the gait pattern.

## AUTHOR CONTRIBUTIONS

AF, FDG, and IJ conceptualized the methods; AF, LP, HK, MW, SVR, HH, EP, and LBO processed the data; AF performed the formal analysis; AF, HK, LBO, AH, KD, GM, AVC, FDG, and IJ acquired funding; AF, FDG, and IJ conducted the investigation; AF and FDG developed the methodology; AH, KD, GM, AVC, FDG, and IJ administrated the project; EP, LBO, AH, KD, GM, and AVC provided resources; AF developed the software; FDG and IJ supervised the project; AF, FDG, and IJ validated the research outputs; AF prepared the data visualization; AF drafted the manuscript; and all authors edited the manuscript.

## DATA AVAILABILITY STATEMENT

All data, code, and materials used in this study will be made available at https://simtk.org/projects/predictcpgait upon publication.

