## Supplementary Material for "Physics-based predictive simulations to explore the differential effects of motor control and musculoskeletal deficits on gait dysfunction in cerebral palsy: a retrospective case study"

### 1 SUPPLEMENTARY TABLES

**Table S1. Generic and personalized optimal fiber lengths (cm).** L and R are for left and right, respectively. Lh and sh are for long and short head, respectively.

| | **Gluteus maximus 1** | | **Gluteus maximus 2** | | **Gluteus maximus 3** | | **Gluteus medius 1** | | **Gluteus medius 2** | |
|---|---|---|---|---|---|---|---|---|---|---|
| | L | R | L | R | L | R | L | R | L | R |
| **Generic** | 13.35 | 13.44 | 12.93 | 12.95 | 12.85 | 12.72 | 6.23 | 6.11 | 10.36 | 10.34 |
| **Personalized** | 6.67 | 6.72 | 6.47 | 6.47 | 7.99 | 7.61 | 3.11 | 3.05 | 5.18 | 5.45 |
| | **Gluteus medius 3** | | **Gluteus minimus 1** | | **Gluteus minimus 2** | | **Gluteus minimus 3** | | **Adductor longus** | |
| | L | R | L | R | L | R | L | R | L | R |
| **Generic** | 6.82 | 6.76 | 6.80 | 6.99 | 6.81 | 6.93 | 3.66 | 3.62 | 13.01 | 12.40 |
| **Personalized** | 3.55 | 3.38 | 4.56 | 4.80 | 3.73 | 3.93 | 2.05 | 1.95 | 6.51 | 6.20 |
| | **Adductor brevis** | | **Adductor magnus 1** | | **Adductor magnus 2** | | **Adductor magnus 3** | | **Pectineus** | |
| | L | R | L | R | L | R | L | R | L | R |
| **Generic** | 13.93 | 13.08 | 9.10 | 8.29 | 11.67 | 11.19 | 12.63 | 13.13 | 16.06 | 14.41 |
| **Personalized** | 8.68 | 8.27 | 7.55 | 7.19 | 5.88 | 5.60 | 6.63 | 6.56 | 10.04 | 9.57 |
| | **Iliacus** | | **Psoas** | | **Quadratus femoris** | | **Gemellus** | | **Piriformis** | |
| | L | R | L | R | L | R | L | R | L | R |
| **Generic** | 10.35 | 10.06 | 9.50 | 9.35 | 3.71 | 3.72 | 1.67 | 1.57 | 2.28 | 2.31 |
| **Personalized** | 5.17 | 5.03 | 4.75 | 4.68 | 4.16 | 3.96 | 1.90 | 1.81 | 1.21 | 1.15 |
| | **Tensor fasciae latae** | | **Gracilis** | | **Semimembranosus** | | **Semitendinosus** | | **Biceps femoris lh** | |
| | L | R | L | R | L | R | L | R | L | R |
| **Generic** | 8.81 | 8.88 | 33.36 | 33.23 | 7.48 | 7.50 | 18.32 | 18.42 | 10.05 | 10.08 |
| **Personalized** | 9.28 | 8.84 | 16.68 | 16.61 | 7.88 | 7.75 | 10.26 | 10.80 | 10.03 | 10.48 |
| | **Biceps femoris sh** | | **Sartorius** | | **Rectus femoris** | | **Vastus medius** | | **Vastus intermedius** | |
| | L | R | L | R | L | R | L | R | L | R |
| **Generic** | 13.54 | 14.08 | 56.42 | 56.73 | 7.74 | 7.73 | 8.09 | 8.02 | 7.66 | 7.59 |
| **Personalized** | 7.39 | 7.04 | 38.74 | 36.89 | 8.16 | 8.59 | 4.93 | 4.70 | 4.76 | 4.73 |
| | **Vastus lateralis** | | **Gastrocnemius medialis** | | **Gastrocnemius lateralis** | | **Soleus** | | **Tibialis posterior** | |
| | L | R | L | R | L | R | L | R | L | R |
| **Generic** | 7.49 | 7.43 | 3.86 | 3.89 | 5.56 | 5.58 | 2.62 | 2.64 | 2.72 | 2.74 |
| **Personalized** | 6.50 | 6.23 | 3.26 | 3.43 | 3.21 | 3.38 | 1.93 | 2.03 | 1.44 | 1.37 |
| | **Tibialis anterior** | | **Extensor digitalis** | | **Extensor hallucis** | | **Flexor digitalis** | | **Flexor hallucis** | |
| | L | R | L | R | L | R | L | R | L | R |
| **Generic** | 8.45 | 8.52 | 8.83 | 8.84 | 9.51 | 9.54 | 2.90 | 2.91 | 3.64 | 3.65 |
| **Personalized** | 4.22 | 4.26 | 4.64 | 4.42 | 5.01 | 4.77 | 1.53 | 1.45 | 1.92 | 1.83 |
| | **Peroneus brevis** | | **Peroneus longus** | | **Peroneus tertius** | | | | | |
| | L | R | L | R | L | R | | | | |
| **Generic** | 4.15 | 4.23 | 4.26 | 4.28 | 6.32 | 6.50 | | | | |
| **Personalized** | 2.19 | 2.12 | 2.24 | 2.14 | 3.35 | 3.25 | | | | |

**Table S2. Generic and personalized tendon slack lengths (cm).** L and R are for left and right, respectively. Lh and sh are for long and short head, respectively.

| | **Gluteus maximus 1** | | **Gluteus maximus 2** | | **Gluteus maximus 3** | | **Gluteus medius 1** | | **Gluteus medius 2** | |
|---|---|---|---|---|---|---|---|---|---|---|
| | L | R | L | R | L | R | L | R | L | R |
| **Generic** | 11.75 | 11.83 | 11.17 | 11.18 | 10.18 | 10.07 | 9.08 | 8.90 | 4.54 | 4.53 |
| **Personalized** | 11.34 | 11.93 | 12.01 | 11.92 | 13.23 | 13.51 | 9.94 | 10.46 | 8.35 | 8.79 |
| | **Gluteus medius 3** | | **Gluteus minimus 1** | | **Gluteus minimus 2** | | **Gluteus minimus 3** | | **Adductor longus** | |
| | L | R | L | R | L | R | L | R | L | R |
| **Generic** | 5.60 | 5.55 | 1.6 | 1.64 | 3.16 | 3.22 | 4.92 | 4.85 | 10.42 | 9.93 |
| **Personalized** | 8.47 | 8.48 | 3.12 | 3.29 | 6.11 | 6.43 | 7.06 | 6.88 | 11.85 | 11.29 |
| | **Adductor brevis** | | **Adductor magnus 1** | | **Adductor magnus 2** | | **Adductor magnus 3** | | **Pectineus** | |
| | L | R | L | R | L | R | L | R | L | R |
| **Generic** | 1.36 | 1.28 | 6.27 | 5.72 | 12.54 | 12.03 | 25.07 | 26.06 | 0.12 | 0.11 |
| **Personalized** | 2.69 | 2.56 | 4.81 | 4.58 | 13.12 | 12.50 | 27.07 | 25.78 | 0.23 | 0.22 |
| | **Iliacus** | | **Psoas** | | **Quadratus femoris** | | **Gemellus** | | **Piriformis** | |
| | L | R | L | R | L | R | L | R | L | R |
| **Generic** | 9.31 | 9.05 | 10.33 | 10.16 | 1.65 | 1.66 | 2.71 | 2.55 | 9.29 | 9.40 |
| **Personalized** | 13.49 | 13.39 | 15.19 | 14.93 | 1.37 | 1.30 | 3.47 | 3.30 | 10.40 | 10.85 |
| | **Tensor fasciae latae** | | **Gracilis** | | **Semimembranosus** | | **Semitendinosus** | | **Biceps femoris lh** | |
| | L | R | L | R | L | R | L | R | L | R |
| **Generic** | 39.41 | 39.73 | 13.27 | 13.22 | 33.56 | 33.64 | 23.88 | 24.01 | 31.45 | 31.53 |
| **Personalized** | 37.94 | 36.14 | 20.25 | 20.21 | 30.10 | 31.68 | 30.11 | 31.69 | 28.75 | 30.26 |
| | **Biceps femoris sh** | | **Sartorius** | | **Rectus femoris** | | **Vastus medius** | | **Vastus intermedius** | |
| | L | R | L | R | L | R | L | R | L | R |
| **Generic** | 7.83 | 8.14 | 3.90 | 3.92 | 31.88 | 31.85 | 11.46 | 11.36 | 11.97 | 11.87 |
| **Personalized** | 10.38 | 10.93 | 7.80 | 7.42 | 28.93 | 29.40 | 14.25 | 14.95 | 14.30 | 15.05 |
| | **Vastus lateralis** | | **Gastrocnemius medialis** | | **Gastrocnemius lateralis** | | **Soleus** | | **Tibialis posterior** | |
| | L | R | L | R | L | R | L | R | L | R |
| **Generic** | 14.01 | 13.88 | 34.99 | 35.23 | 33.44 | 33.56 | 23.41 | 23.61 | 27.22 | 27.35 |
| **Personalized** | 14.13 | 14.87 | 33.23 | 34.59 | 33.38 | 34.48 | 23.53 | 23.52 | 29.27 | 29.16 |
| | **Tibialis anterior** | | **Extensor digitalis** | | **Extensor hallucis** | | **Flexor digitalis** | | **Flexor hallucis** | |
| | L | R | L | R | L | R | L | R | L | R |
| **Generic** | 19.23 | 19.39 | 29.86 | 29.91 | 26.12 | 26.21 | 34.07 | 34.18 | 32.18 | 32.3 |
| **Personalized** | 20.40 | 21.48 | 30.93 | 31.75 | 27.67 | 28.43 | 35.60 | 35.49 | 34.12 | 33.87 |
| | **Peroneus brevis** | | **Peroneus longus** | | **Peroneus tertius** | | | | | |
| | L | R | L | R | L | R | | | | |
| **Generic** | 13.37 | 13.63 | 30.01 | 30.11 | 8.00 | 8.23 | | | | |
| **Personalized** | 15.09 | 15.88 | 31.25 | 31.40 | 9.66 | 10.17 | | | | |

### 2 SUPPLEMENTARY FIGURES

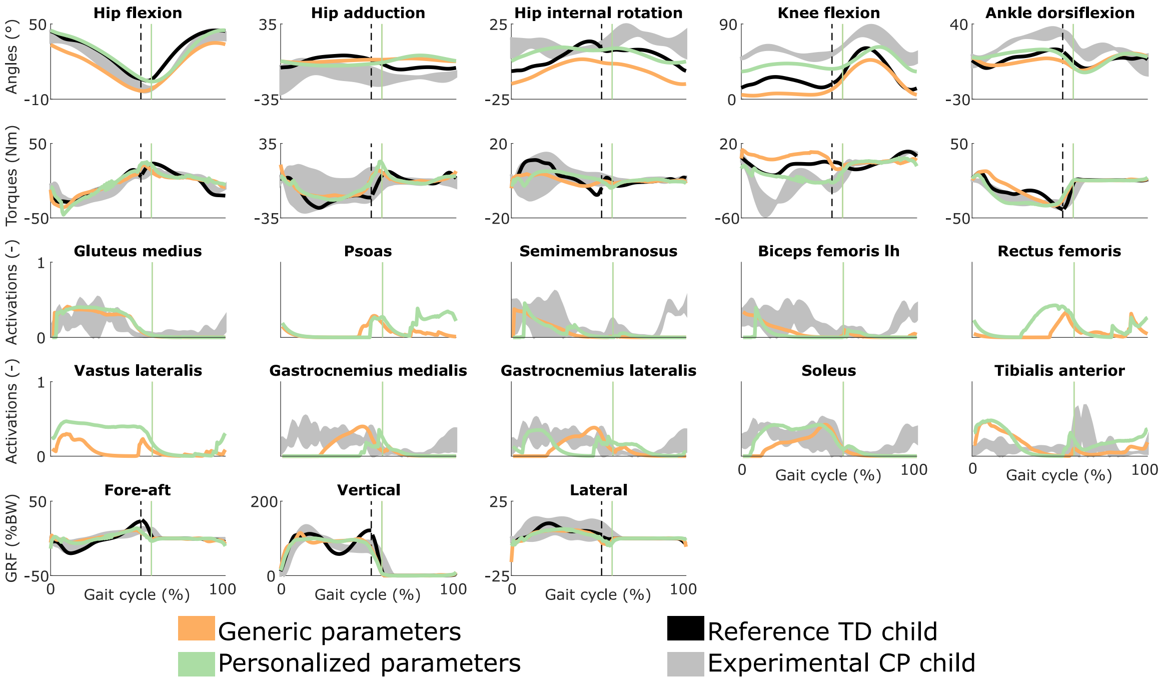


**Figure S1. Influence of the muscle-tendon parameters on the predicted walking gaits.** Variables from the left leg are shown over a complete gait cycle; right leg variables are shown in Figure 2 (Manuscript). Solid vertical lines indicate the transition from stance to swing. Experimental data is shown as mean ± two standard deviations. Experimental EMG data was normalized to peak activations. Reference TD child data was available for a single gait cycle starting at right heel strike; left leg data was thus reconstructed from that gait cycle but is discontinuous as indicated by the dashed vertical lines. GRF is for ground reaction forces; BW is for body weight; COT is for metabolic cost of transport; lh is for long head.

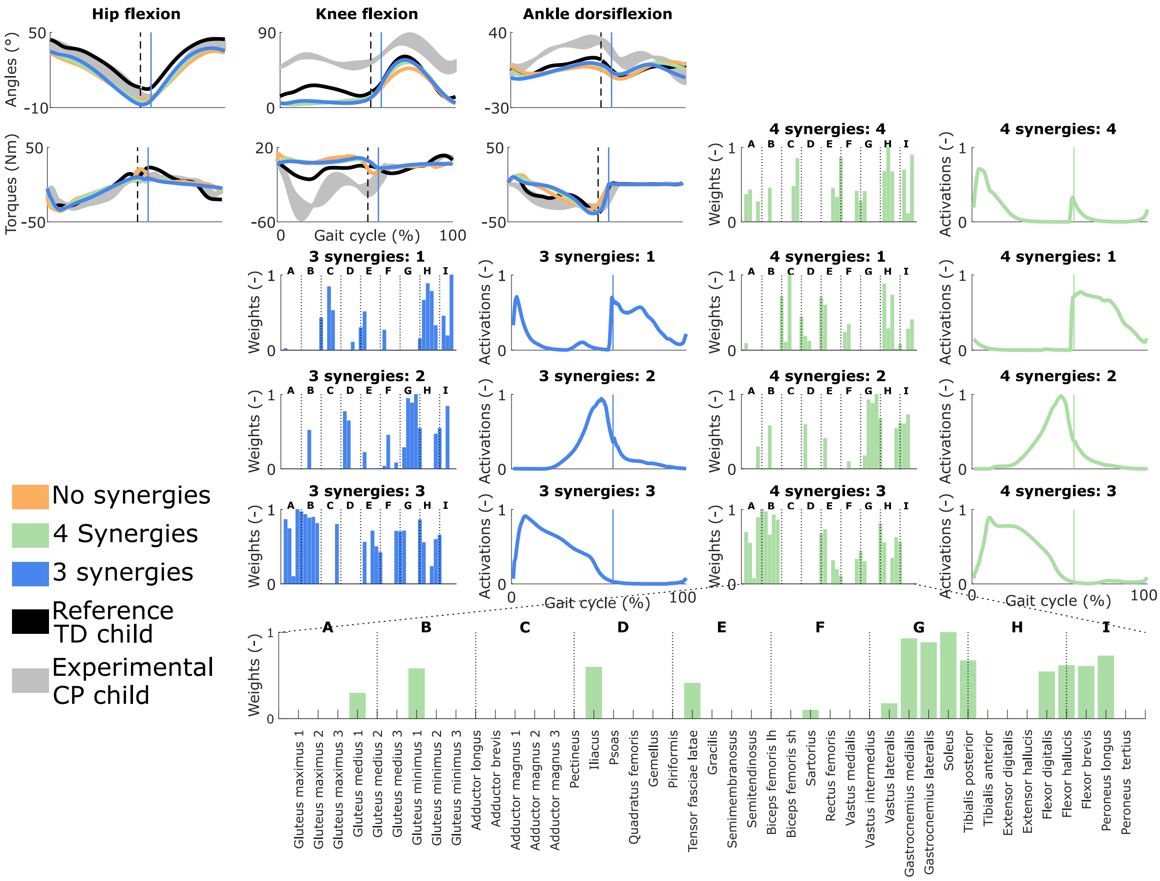


**Figure S2. Influence of the synergies on walking gaits predicted with the generic muscle-tendon parameters.** Variables from the left leg are shown over a complete gait cycle; right leg variables are shown in Figure 3 (Manuscript). Vertical lines (solid) indicate the transition from stance to swing. Panels of synergy weights are divided into sections (A-I) to relate bars to muscle names provided in the bottom bar plot, which is an expanded version of the plot of weights with title 4 synergies: 3. Lh and sh are for long and short head, respectively. Weights were normalized to one. Experimental data is shown as mean ± two standard deviations. Reference TD child data was available for a single gait cycle starting at right heel strike; left leg data was thus reconstructed from that gait cycle but is discontinuous as indicated by the dashed vertical lines.

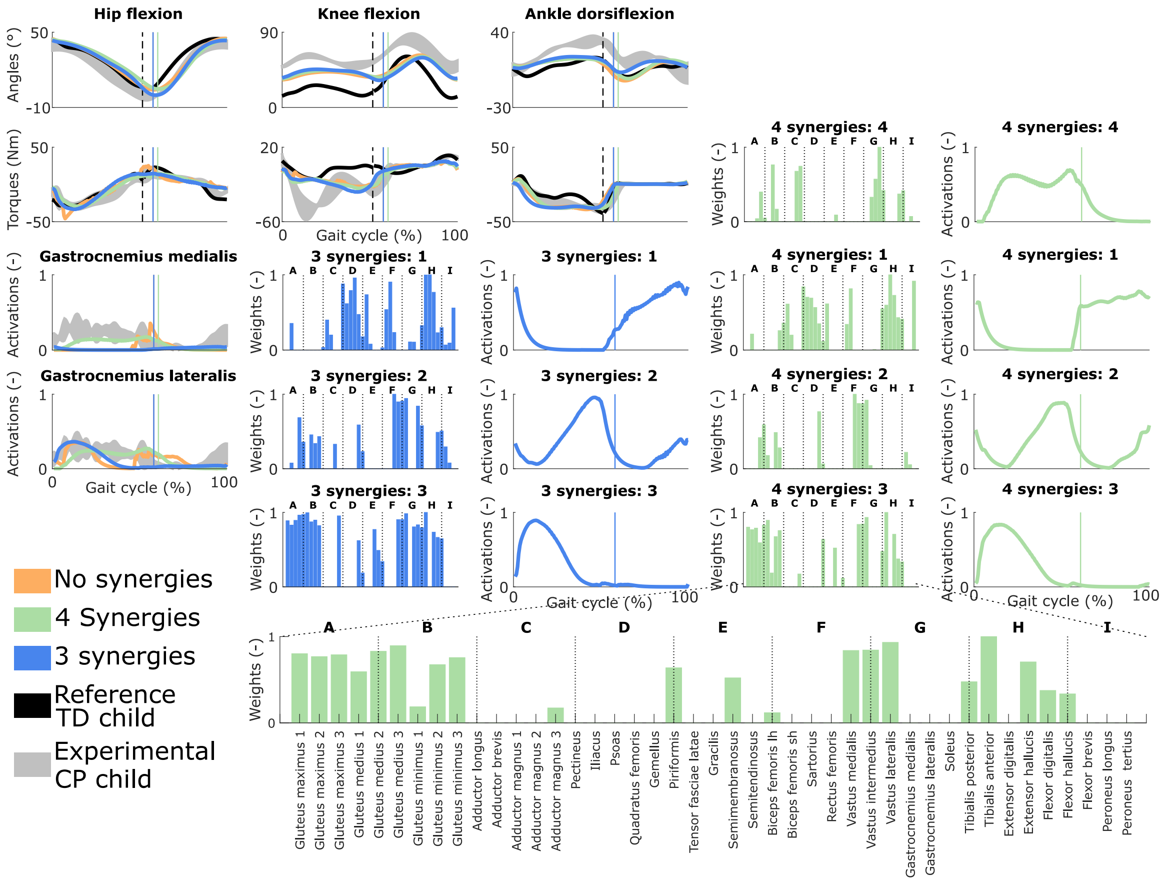


**Figure S3. Influence of the synergies on walking gaits predicted with the personalized muscle-tendon parameters.** Variables from the left leg are shown over a complete gait cycle; right leg variables are shown in Figure 4 (Manuscript). Vertical lines (solid) indicate the transition from stance to swing. Panels of synergy weights are divided into sections (A-I) to relate bars to muscle names provided in the bottom bar plot, which is an expanded version of the plot of weights with title 4 synergies: 3. Lh and sh are for long and short head, respectively. Weights were normalized to one. Experimental data is shown as mean ± two standard deviations. Experimental EMG data was normalized to peak activations. Reference TD child data was available for a single gait cycle starting at right heel strike; left leg data was thus reconstructed from that gait cycle but is discontinuous as indicated by the dashed vertical lines.

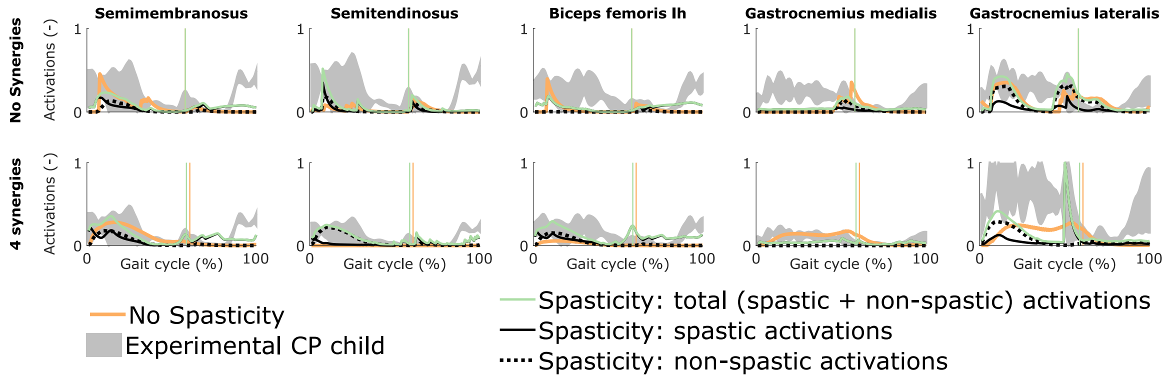


**Figure S4. Influence of spasticity on the predicted muscle activity**. Activations from left leg muscles only are shown over a complete gait cycle; right leg activations are shown in Figure 5 (Manuscript). When accounting for spasticity, total activations (green) combine spastic (solid black) and non-spastic (dotted black) activations. Vertical lines indicate the transition from stance to swing. Experimental data is shown as mean ± two standard deviations. Experimental EMG data was normalized to peak activations. Lh is for long head.

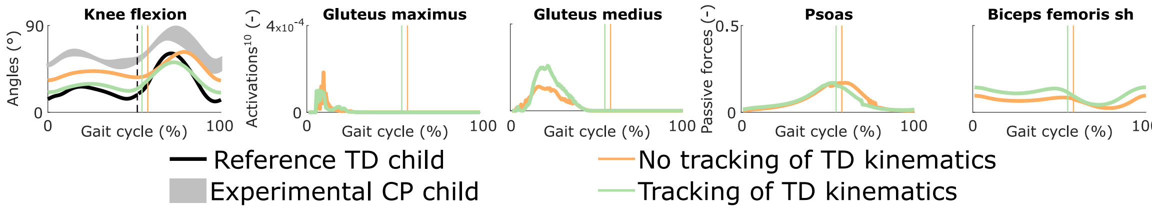


**Figure S5. Influence of tracking the TD kinematics on predicted walking gaits.** Variables from the left leg are shown over a complete gait cycle; right leg variables are shown in Figure 6 (Manuscript). Vertical lines indicate the transition from stance to swing. Experimental data is shown as mean ± two standard deviations. Muscle fatigue is modeled by activations at the tenth power. Passive muscle forces are normalized by maximal isometric muscle forces. Sh is for short head.

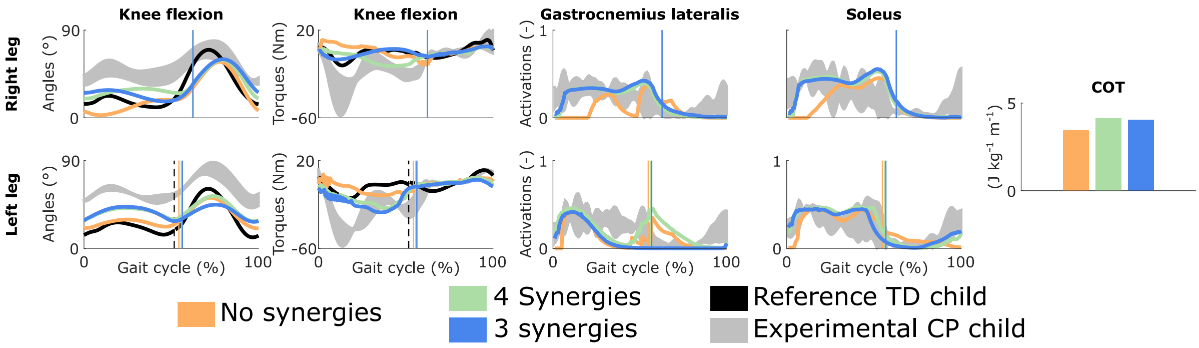


**Figure S6. Influence of the synergies on walking gaits predicted with the personalized muscle-tendon parameters while tracking the TD kinematics.** Solid vertical lines indicate the transition from stance to swing. Experimental data is shown as mean ± two standard deviations. Experimental EMG data was normalized to peak activations. Reference TD child data was available for a single gait cycle starting at right heel strike; left leg data was thus reconstructed from that gait cycle but is discontinuous as indicated by the dashed vertical lines.
